## Supplementary figures and images for "SARS-CoV-2 sequencing artifacts associated with targeted PCR enrichment and read mapping"

### Figure S1

#### Base calls generated for samples sequenced with Artic V3 and V4.1

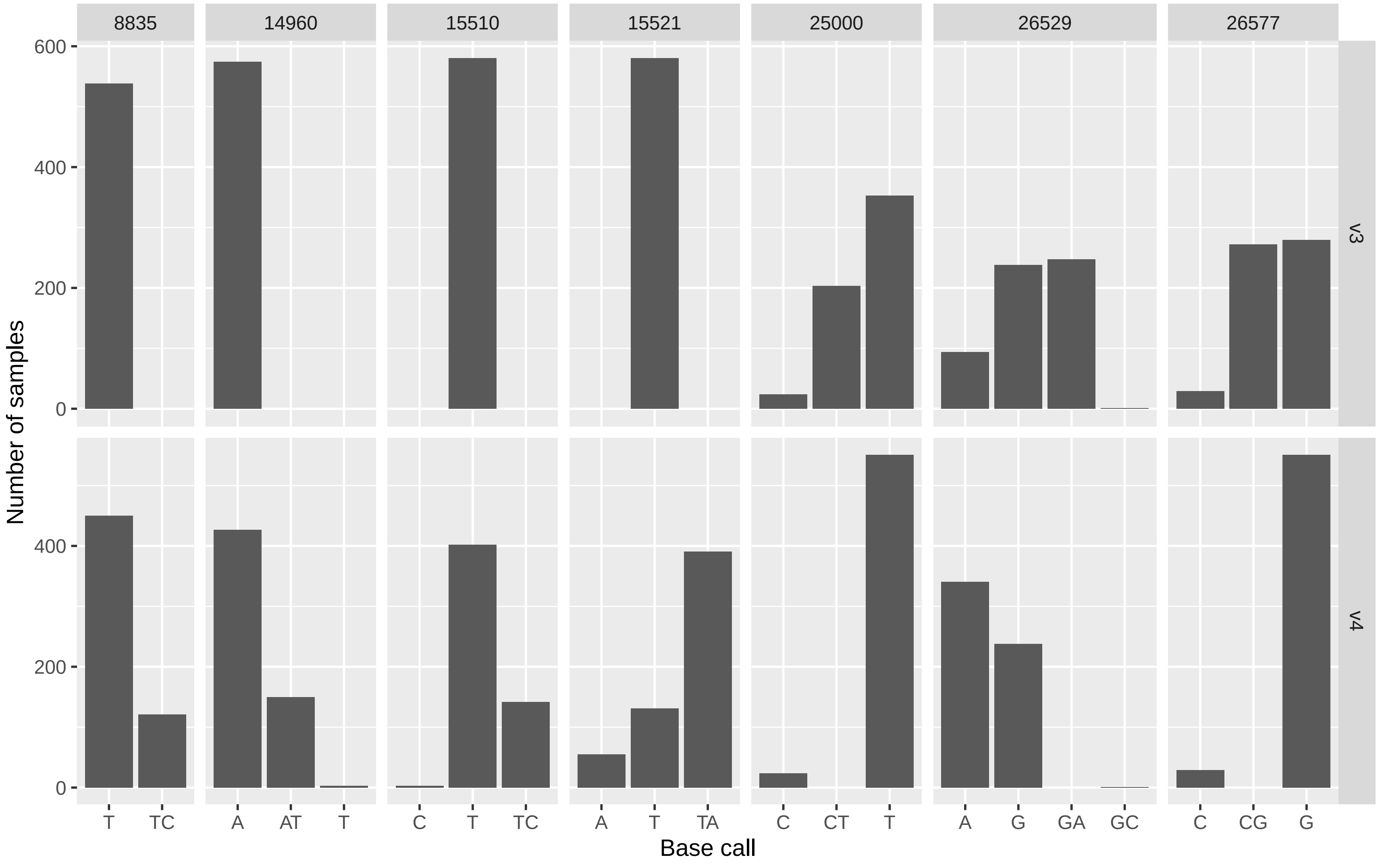

### Figure S2

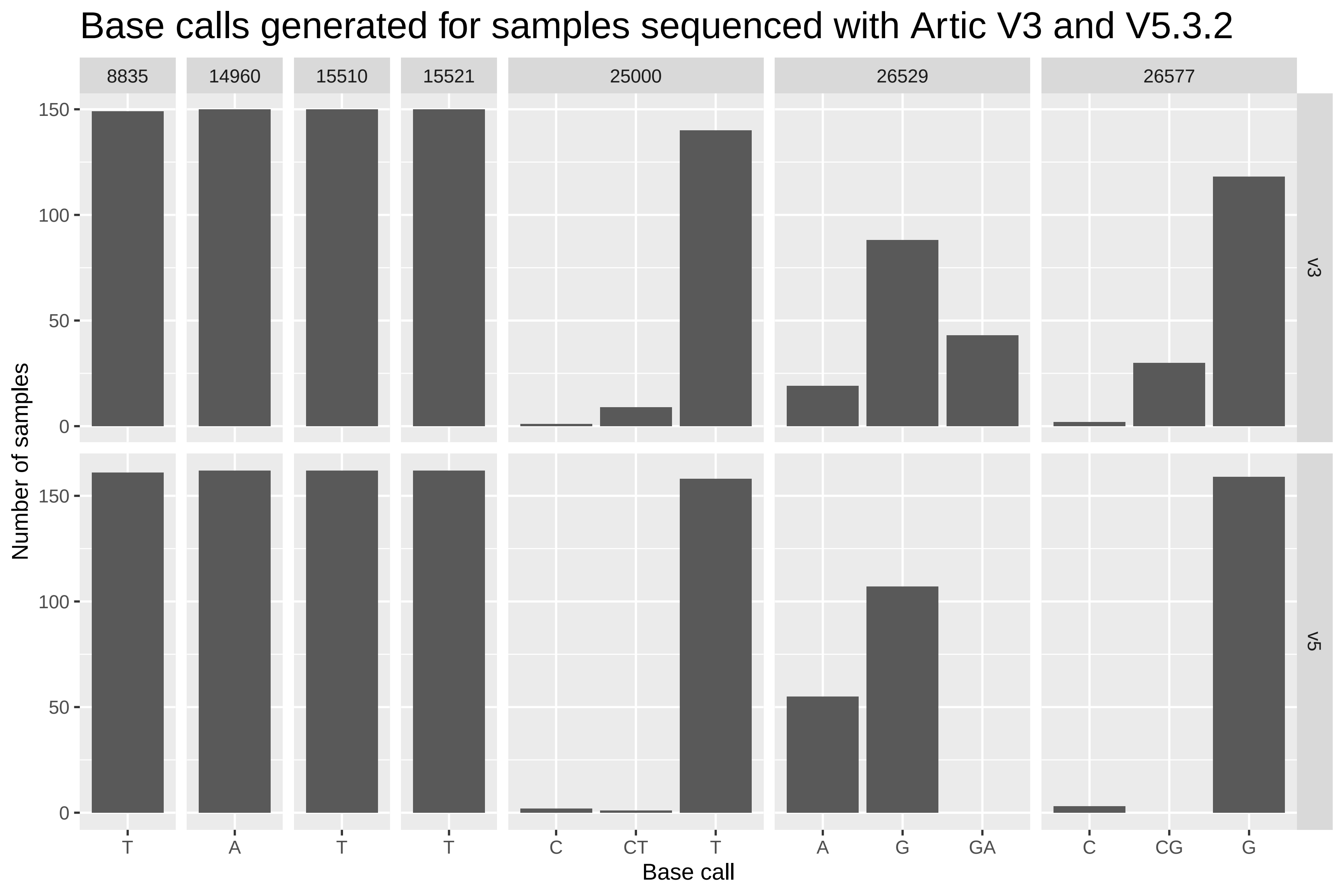
